## Supplemental Figures S1-S10 for "Differential contribution of two organelles of endosymbiotic origin to iron-sulfur cluster synthesis and overall fitness in Toxoplasma"

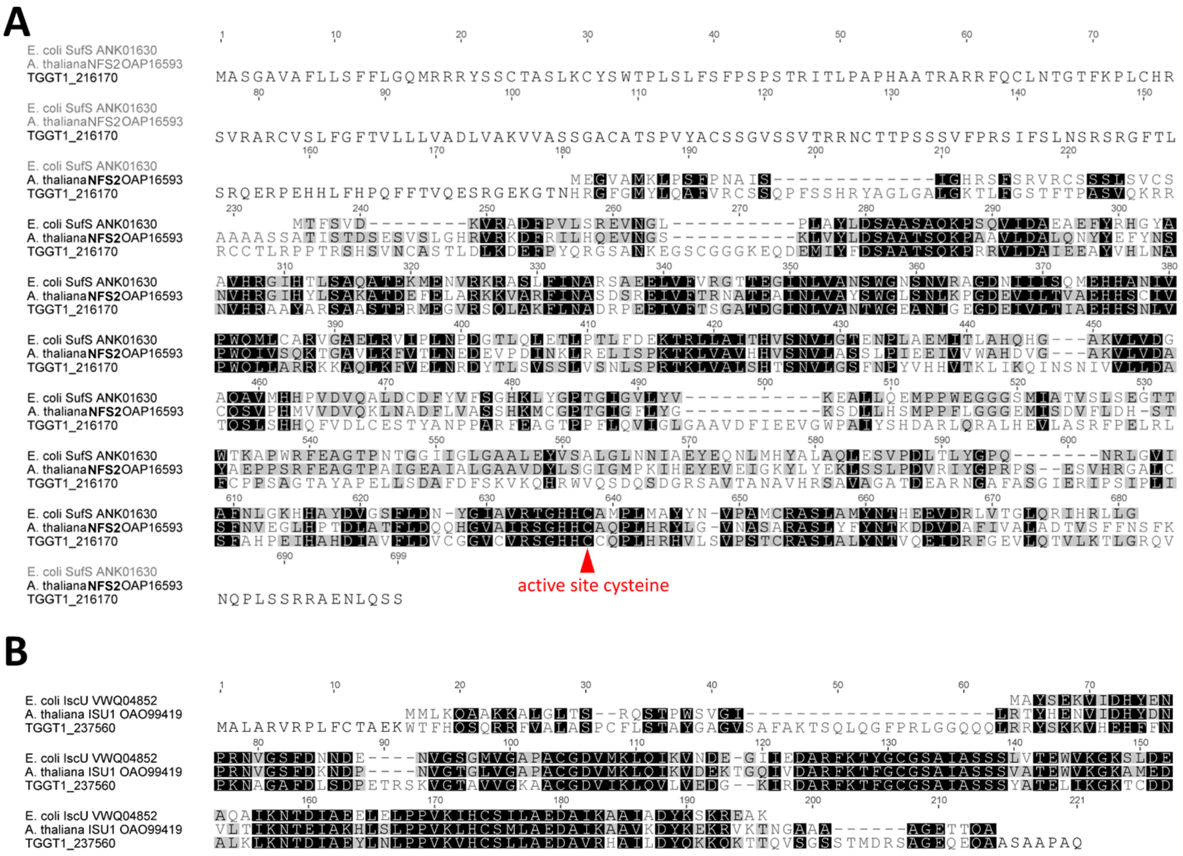

**Figure S1. Alignment of SufS/NFS2 and IscU/ISU1 homologs**. TgNFS2 (A) and TgISU1 (B) homologs were aligned to their counterparts from plant (*Arabidopsis thaliana*) and bacteria (*Escherishia coli*). Key conserved cysteine residue for cysteine desulfurase activity is indicated.

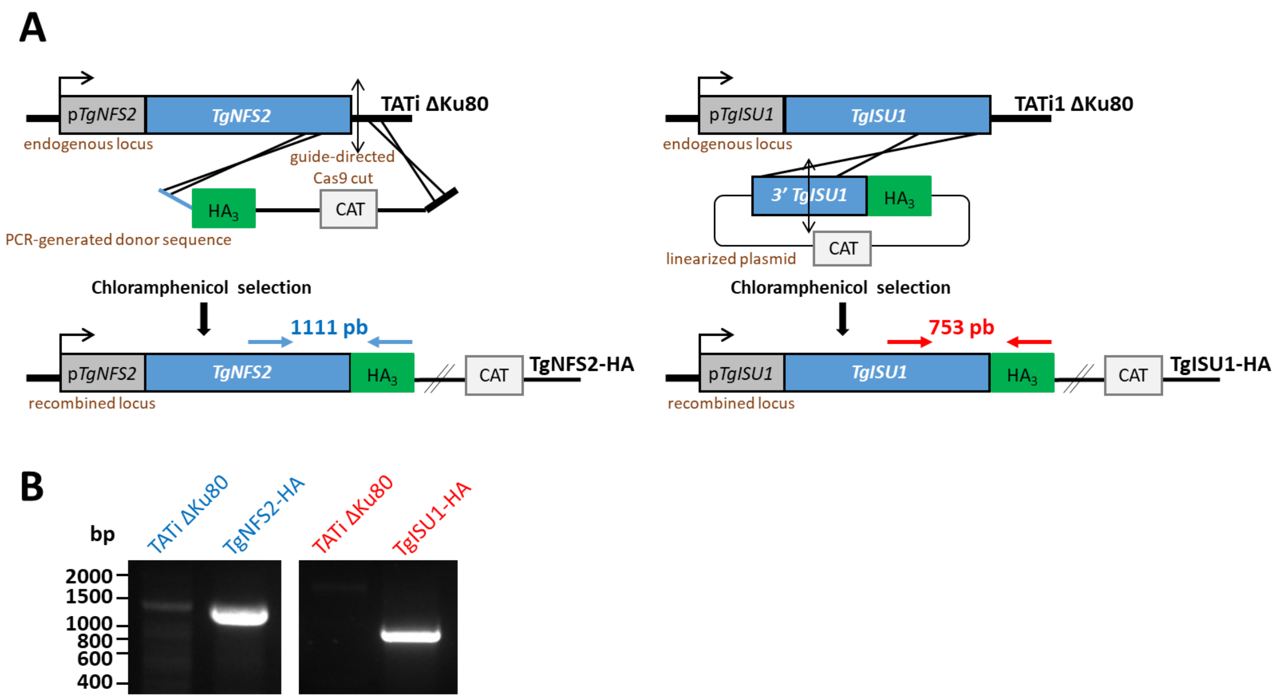


**Figure S2. Generation of HA-tagged TgNFS2 and TgISU1 cell lines.** A) Schematic representation of the strategy for expressing HA-tagged versions of TgNFS2 (left) and TgISU1 (right) by homologous recombination at the native locus of the corresponding gene of interest . Chloramphenicol was used to select transgenic parasites based on their expression of the Chloramphenicol acetyltransferase (CAT). B) Diagnostic PCR for verifying correct integration of the construct. The amplified fragments correspond to the blue or red arrows in A), and specific primers used were ML3982/ML1476 (TgNFS2) and ML4208/ML1476 (TgISU1).

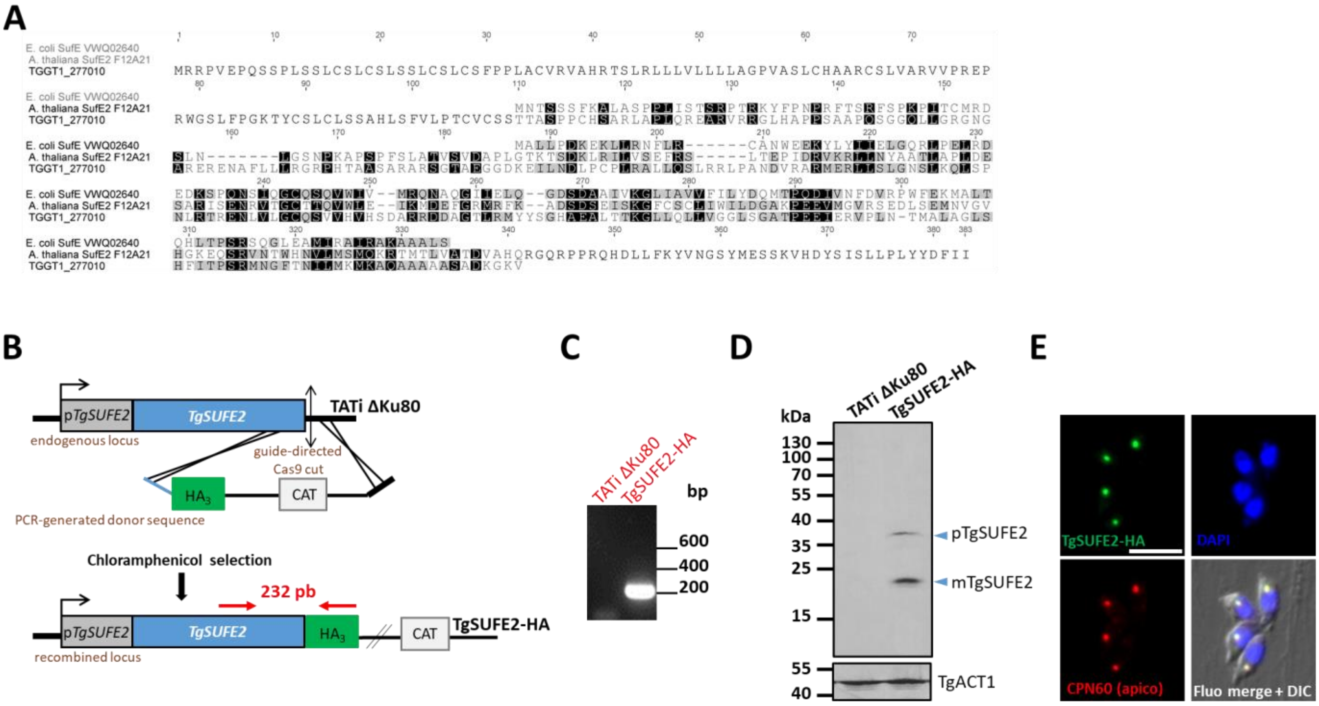


**Figure S3. HA-tagging of TgSUFE2 shows it is an apicoplast protein.** A) Sequence alignment of TgSUFE2 (TGGT1_277010) with plant (*A. thaliana*) and bacterial (*E. coli*) homologues. B) Schematic representation of the strategy for expressing an HA-tagged version of TgSUFE2 by double homologous recombination at the native locus. Chloramphenicol was used to select transgenic parasites based on their expression of the Chloramphenicol acetyltransferase (CAT). C) Diagnostic PCR for verifying correct integration of the construct. The amplified fragment corresponds to the red arrows in B), and specific primers used were ML4101/ML1476. D) Detection by immunoblot of C-terminally HA-tagged TgSUFE2 in parasite extracts reveals the presence of both precusor and mature forms of the protein. Anti-actin antibody (TgACT1) was used as a loading control. E) Immunofluorescence assay shows TgSUFE2 co-localizes with apicoplast marker TgCPN60. Scale bar represents 5 µm. DNA was labelled with DAPI. DIC: differential interference contrast.

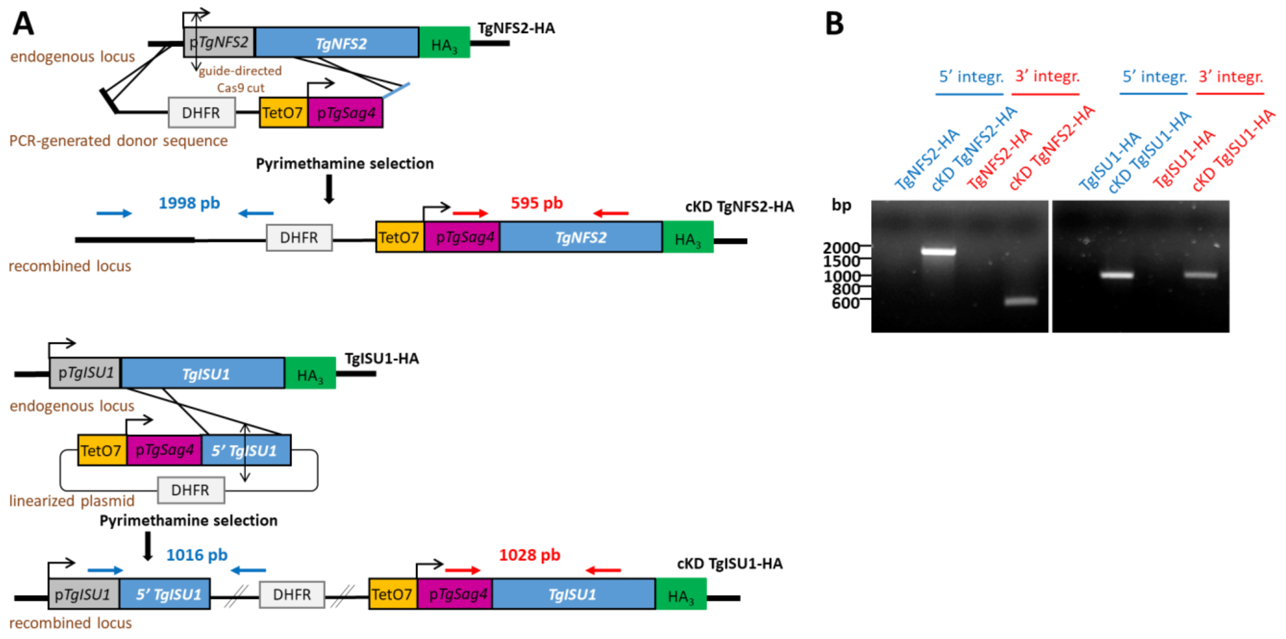


**Figure S4. Generation of TgNFS2 and TgISU1 conditional mutants.** A) Schematic representation of the strategy for generating TgNFS2 (top) and TgISU1 (bottom) conditional knock-down cell lines by homologous recombination at the native locus. Pyrimethamine was used to select transgenic parasites based on their expression of Dihydrofolate reductase (DHFR). B) Diagnostic PCR for verifying correct integration of the construct. The amplified fragments confirming 5' and 3' integration correspond to the blue and red arrows displayed in A), respectively, and specific primers used were: ML4158/ML687 (TgNFS2 5' integration), ML1041/ML4159 (TgNFS2 3' integration), ML1774/ML4388 (TgISU1 5' integration), ML1771/ML4387 (TgISU1 3' integration).

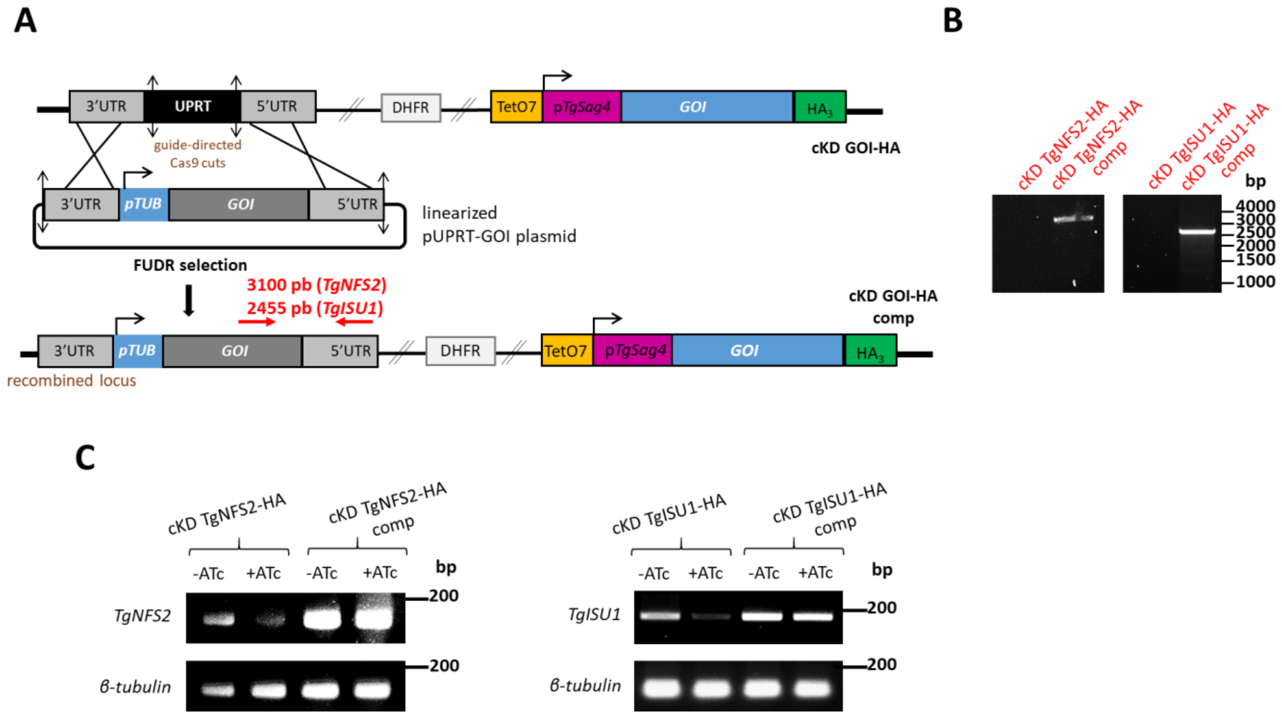


**Figure S5. Generation of TgNFS2 and TgISU1 complemented cell lines.** A) Schematic representation of the strategy for generating TgNFS2 and TgISU1 complemented cell lines by integrating an extra copy of the gene of interest (GOI) by double homologous recombination at the *Uracil Phosphoribosyltransferase* (*UPRT*) locus. Negative selection with 5-fluorodeoxyuridine (FUDR) was used to select transgenic parasites based on their absence of UPRT expression. B) Diagnostic PCR for verifying correct integration of the construct. The amplified fragments confirming integration correspond to the red arrows displayed in A), and specific primers used were: ML2866/ML4686 (TgNFS2 integration), ML2866/ML4455 (TgISU1 5' integration). C) Semi-quantitative RT-PCR analysis from cKD TgNFS2-HA, cKD TgISU1-HA and their respective complemented cell lines grown for three days in the presence or absence of ATc, using specific primers couples ML4686/ML4687 *(TgNFS2)* and ML4684/ML4685 (*TgISU1*). It shows complemented cell lines express high levels of the corresponding mRNA. Specific *β-tubulin* primers (ML841/ML842) were used as controls.

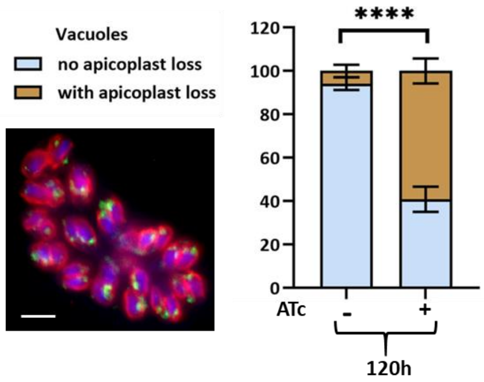


**Figure S6. Quantification of apicoplast loss upon TgNFS2 depletion using streptavidin.** Percentage of cKD TgNFS2-HA parasites-containing vacuoles displaying a loss of apicoplast signal when labeled with streptavidin-Fluorescein Isothiocyanate after culture in the presence or absence of ATc for 120 hours. Data are mean values from $n$=3 independent experiments ±SEM. **** $p \leq 0.0001$, Student's $t$-test. Inset : typical streptavidin labeling of the apicoplast (green) in a tachyzoites-containing vacuole, the inner membrane complex was stained with anti-TgIMC3 (red) and the DNA with DAPI (blue); scale bar=5µm.

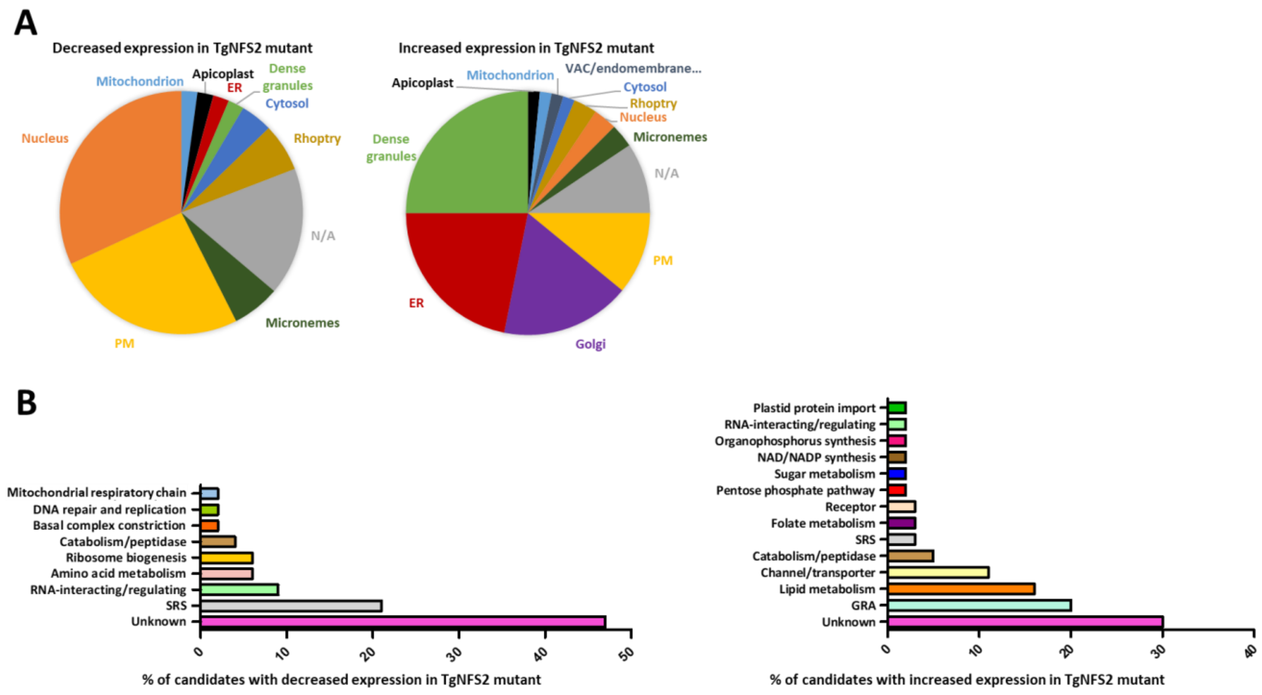


**Figure S7. Quantitative proteomics shows depletion of TgNFS2 does not have a global impact on the apicoplast, but may suggest compensatory response from other cellular pathways in response to specific apicoplast-related lipid synthesis defects.** Classification of variant proteins according to their putative cellular localization (A) and function (B). N/A: not available; ER: endoplasmic reticulum; PM: plasma membrane; VAC: vacuolar compartment; GRA: dense granule protein; SRS: SAG-related sequence. In particular, the increased expression of ER-located lipid metabolism enzymes suggests possible compensation for loss of apicoplast-related lipid synthesis function.

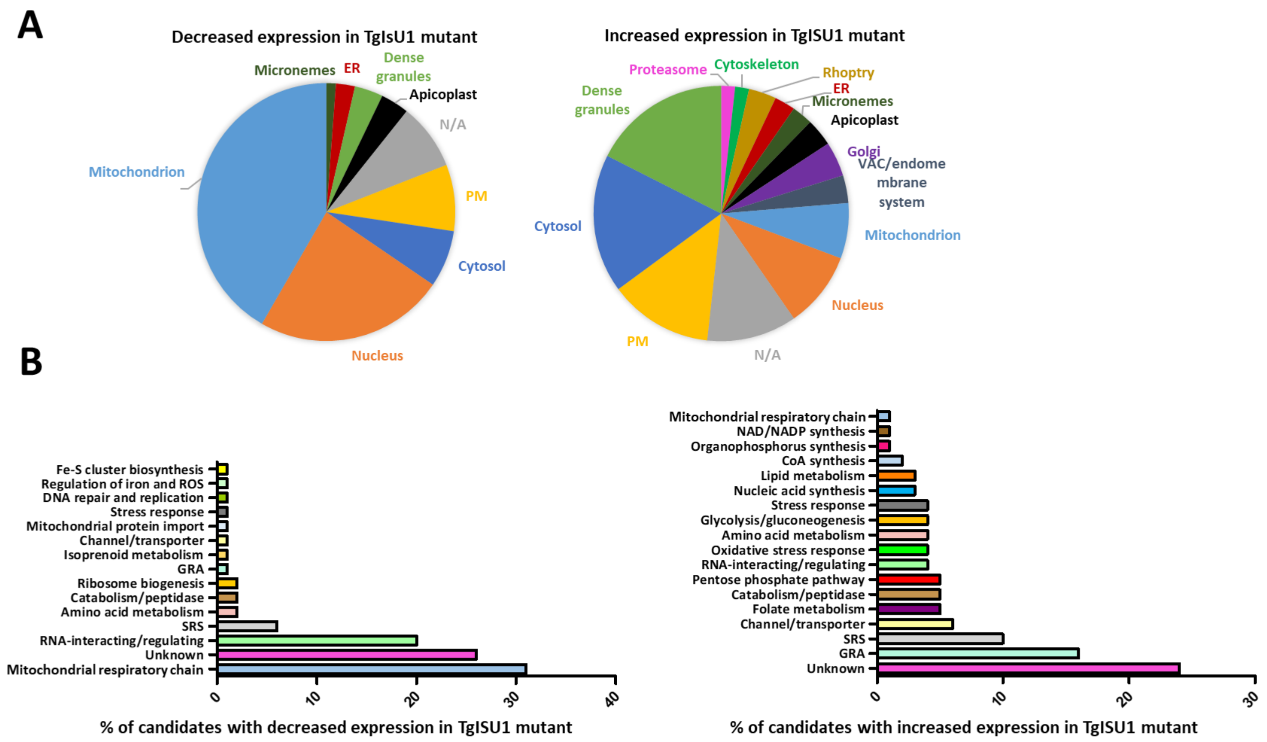


**Figure S8. TgISU1-depleted parasites show a marked decrease in proteins related to mitochondrial respiration, and a strong increase in bradyzoite-specific dense granules proteins and surface antigens.** Classification of variant proteins according to their putative cellular localization (A) and function (B). N/A: not available; ER: endoplasmic reticulum; PM: plasma membrane; VAC: vacuolar compartment; GRA: dense granule protein; SRS: SAG-related sequence. A large proportion of components of complexes II, III and IV of the mitochondrial respiratory chain, which involve Fe-S proteins, were found to be less abundant. Conversely, the abundance of many bradyzoite-specific dense granule proteins of plasma membrane-located surface antigens increased.

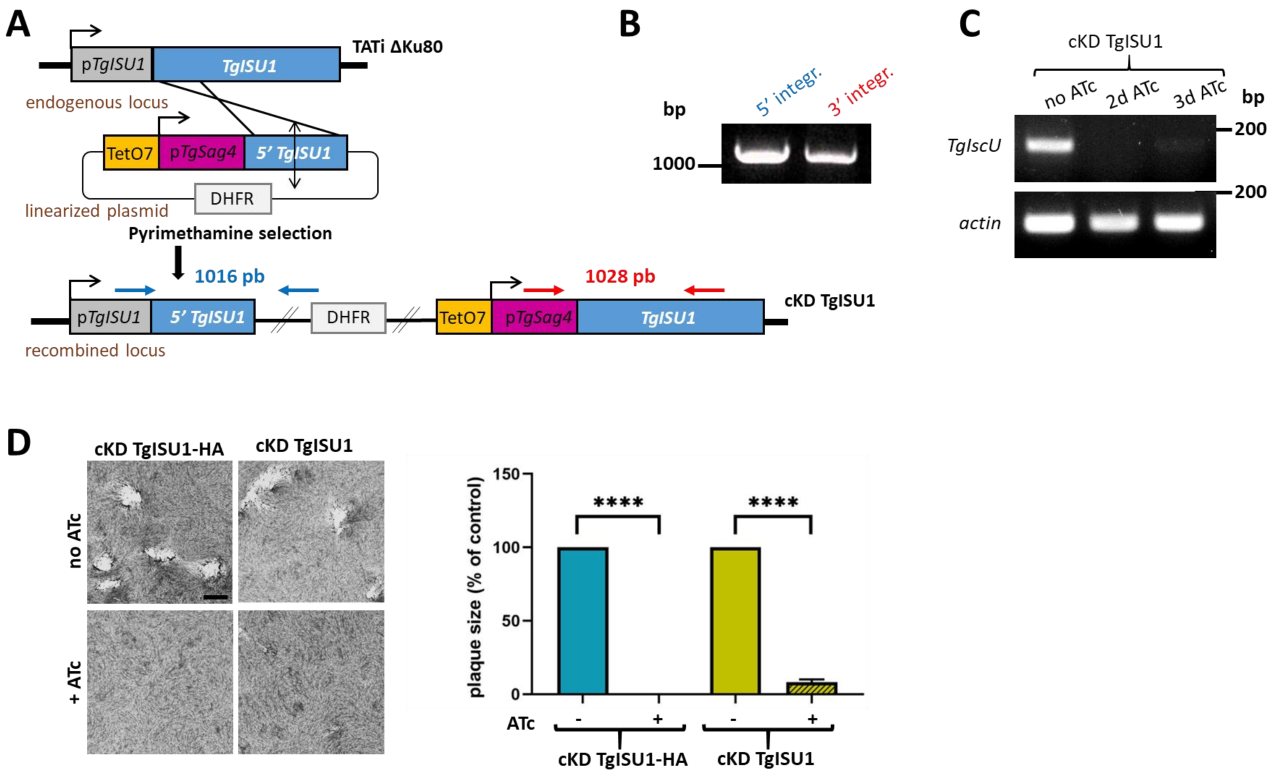


**Figure S9. Generation of a tag-free cKD TgISU1 cell line.** A) Schematic representation of the strategy for generating the conditional knock-down cell line by homologous recombination at the native locus. Pyrimethamine was used to select transgenic parasites based on their expression of Dihydrofolate reductase (DHFR). B) Diagnostic PCR for verifying correct integration of the construct. The amplified fragments confirming 5' and 3' integration correspond to the blue and red arrows displayed in A), respectively, and specific primers used were ML1774/ML4388 (5' integration), and ML1771/ML4387 (3' integration). C) Semi-quantitative RT-PCR analysis of the cKD TgISU1 cell line grown for up to three days in the presence or absence of ATc, using specific primers couple ML4684/ML4685, showing effcient down-regulation of *TgISU1* expression. Specific *actin* primers (ML843/ML844) were used as controls. D) Plaque assays were carried out by infecting HFF monolayers with the newly generated cKD TgISU1 cell line or the original cKD TgISU1-HA mutant cell line as a control. They were grown for 7 days ± ATc. Measurements of lysis plaque areas are shown on the right and confirm a significant defect in the lytic cycle in the two mutant cell lines upon ATc addition. Values are means of n=3 experiments ± SEM. **** denotes $p \leq 0.0001$, ANOVA. Scale bar= 1mm.

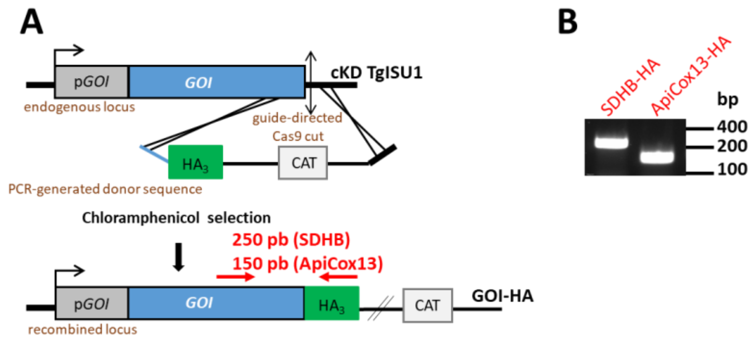


**Figure S10. Tagging of mitochondrial candidates in the cKD TgISU1 background.** A) Schematic representation of the strategy for expressing HA-tagged versions of TgSDHB and TgApiCox13 by homologous recombination at the native locus of the corresponding gene of interest. Chloramphenicol was used to select transgenic parasites based on their expression of the Chloramphenicol acetyltransferase (CAT). B) Diagnostic PCR for verifying correct integration of the construct. The amplified fragment corresponds to the red arrows in A), and specific primers used were ML1476/ML5013 (TgSDHB), ML1476/ML5006 (TgApiCox13).
